# B chromosome and its non-Mendelian inheritance in *Atractylodes lancea*

**DOI:** 10.1101/2024.05.27.595828

**Authors:** Kazuya Hara, Shinji Kikuchi, Misaki Inoue, Takahiro Tsusaka, Miki Sakurai, Hideyuki Tanabe, Kenta Shirasawa, Sachiko Isobe

## Abstract

Supernumerary B chromosomes contribute to intraspecific karyotypic variation. B chromosomes have been detected in more than 2000 organisms; they possess unique and diverse features, including non-Mendelian inheritance. Here we report one or more B chromosomes in the gynodioecious plant, *Atractylodes lancea*. Among 54 *A. lancea* lines, 0–2 B chromosomes were detected in both hermaphroditic and female plants, with the B chromosomes appearing as DAPI-bright regions within the nuclei. Genomic *in situ* hybridization revealed that the B chromosomes had no conserved A chromosome DNA sequences, which was confirmed by fluorescence *in situ* hybridization probed with independently dissected B chromosomes. In male meiosis, the B chromosome did not pair with an A chromosome and was therefore eliminated; accordingly, only 20.1% and 18.6% of these univalent B chromosomes remained at the end of meiosis for the 1B lines of KY17-148 and KY17-118, respectively. However, we also found that B chromosomes were transmitted from male parents in 40.8%–44.2% and 47.2% of the next generation; although these transmission rates from male parents were not essentially different from Mendelian inheritance (0.5), the transmission of gametes carrying B chromosomes increased through fertilization or seed development. B chromosomes were transmitted from three of four 1B female parents to 64.3%–92.6% of the next generation, suggesting B chromosome accumulation. We propose that the B chromosome of *A. lancea* has a specific sequence and persists via non-Mendelian inheritance from female parents. Overall, *A. lancea* is a useful material for understanding the structure, evolution, and mechanism of non-Mendelian inheritance of B chromosomes.

## Introduction

Although each species possesses a particular karyotype, karyotypic changes are occasionally detected within many species. These changes can occur due to polyploidy, aneuploidy, and chromosomal rearrangement events such as chromosomal fusion, fission, and translocation. Moreover, heteromorphic sex chromosomes and XO and ZO sex-determination systems can cause different karyotypes within a species. Since the discovery of supernumerary chromosomes in Hemiptera by Wilson [1], B chromosomes (Bs)—extra dispensable chromosomes—have been detected in more than 2000 plant and animal species [2–5]. Supernumerary B chromosomes were named in contrast to the standard A chromosomes [6], and they also cause karyotypic changes. Because B chromosomes are not essential, individuals with multiple copies (in some case >10) of the B chromosome have been identified within a species [7–8]. In some case, such accessory chromosomes are maintained within a population by non-Mendelian inheritance [9]. The phenomenon of a chromosome being transmitted to progeny at a rate greater than the Mendelian rate (0.5) is known as genetic drive, and this appears in several species as a feature of B chromosomes (Bs); however Bs lacking genetic drive are also known to exist [10]. Genetic drive can take several forms, including nondisjunction, mitotic drive (e.g., in *Crepis capillaris*; [10]), meiotic drive, and transmission drive (or preferential fertilization), before, during, and/or after meiosis [11]. The frequency of B chromosomes within a population is determined by the fitness of the host and the efficiency of the drive mechanism [4].

The drive mechanism of B chromosomes has been thoroughly investigated in rye and maize. In rye, B chromosomes are transmitted into the generative nucleus due to the failure of chromatids to separate during cell division, i.e., via nondisjunction by the heterochromatic end of the long arm and asymmetric spindle formation during pollen (male) mitosis I [3, 12–14]. Interestingly, nondisjunction of rye B chromosomes has also been observed during the first mitosis within the embryo sac (female) [3, 15]. In maize, two dive mechanisms have been reported as linked to the formation of B chromosomes in maize [16], in, (i) nondisjunction during the second pollen mitosis in the generative nucleus [17–19] and (ii) preferential fertilization of the egg cell rather than the central cell by sperm containing two B chromosomes [20]. Beyond rye and maize, in *Lilium callosum*, a female meiotic drive has been detected, wherein the B chromosome passes toward the micropylar side at a frequency of 63.7% [21]. The number of B chromosomes may also differ among organs, for example due to programmed elimination in the roots during embryo development in *Aegilops speltoides* [22].

What are the general structural features of B chromosomes? Although B chromosomes accumulate multiple mutations and structural changes, they are believed to have originated from the A chromosomes of the same species or from a relic of a wide cross such that Bs were introgressed via interspecific cross, after which they persisted and evolved (reviewed by [3]). In plants, B chromosomes are generally smaller than A chromosomes. Structural polymorphisms of B chromosomes, such as large B and micro B, have been reported in the daisy *Brachycome* [23, 24]. Plant B chromosomes may be euchromatic [10], but they are often heterochromatic. This heterochromatic nature derives from the accumulation of several specific repetitive sequences, such as rye D1100, E3900, and ScCI11 [14]. The structural and epigenetic specificity of the B chromosome suppresses its ability to pair with an A chromosome during meiosis, a behaviour that is characteristic of the B chromosomes. Recently, a high-quality sequence of the maize B chromosome was released [25], in which the authors predicted 758 protein-encoding genes (including at least 88 expressed genes) present in 125.9 Mbp of the B chromosome sequence; the absence of synteny among predicted B genes suggested that it evolved via translocation from the A chromosome [25]. Moreover, genes present on the independently evolved B chromosome were found to affect gene expression on the A chromosome [26, 27]. Traits are also present on B chromosomes in other species: for example, the rye B chromosome confers heat tolerance during microsporogenesis [28], and the *Allium* B chromosome promotes rapid seed germination [29, 30].

The gynodioecious species *Atractylodes lancea* De Candolle, in the family Asteraceae (Compositae), is distributed in East Asia [31], and its dried rhizomes have been used as crude drugs, generally to treat digestive disorders and body fluid imbalances, in Chinese and Japanese traditional herbal medicines [32, 33]. Its chromosome number is reported as 2*n* = 24 [34, 35], and genome sequencing, genetic marker development for medicinal compound identification, genetic analysis, and molecular breeding research projects are underway on the basis of this chromosome number [36–38]. However, in this study we report the identification of a novel extra chromosome in *A. lancea.* Our cytological observations revealed that extra chromosomes are widely present in the species. We therefore sought to determine in this study (i) whether this excess chromosome is responsible for karyotypic variation in *A. lancea*, (ii) whether this excess chromosome can be considered a B chromosome insofar as it shows characteristic B chromosome structural features and behaviors, and (iii) whether this extra chromosome causes genetic drive. Finally, we propose *A. lancea* as a new model system for research aiming to understand the structure, evolution, and mechanism of non-Mendelian inheritance mechanisms of B chromosomes.

## Materials and Methods

### Plant materials

*A. lancea* De Candolle was grown in an experimental field at Tsumura and Co., Japan, in 2020, 2021, and 2023. Overall, we used 54 resource lines (see **S1 Table**) of *A. lancea*. Each of these originated from China and was preserved by Tsumura and Co. [37]. Several plants used to generate F_1_ seedlings were prepared in 15 cm pots in a growth chamber with a fixed temperature of 28°C.

### Chromosome preparation

Mitotic chromosomes were obtained from elongated fresh roots of plants grown in 15-cm pots. After excision, roots were treated with 2 mM 8-hydroxyquinoline at 18°C for 6 h then fixed with 3:1 (v/v) ethanol–acetic acid at 18°C for 5 days. Next, fixed roots were stored in 70% ethanol at 4°C until use. Meiotic chromosomes were obtained from young anthers (i.e., 1.0–1.5 mm in length). Excised anthers were fixed using the same procedure used for the root samples.

Next, mitotic and meiotic chromosome slides were prepared using the enzymatic maceration–squash method [39]. The chromosomes were counterstained with 5 μg/mL of 4,6-diamidino-2-phenylindole (DAPI) in Vectashield (Vector Laboratories, USA) then observed under a BX43 phase-contrast microscope or a BX-53 fluorescence microscope (both from Olympus, Japan). All fluorescence images were captured using a CoolSNAP MYO CCD camera (Photometrics, USA) and the resulting images were processed with MetaVue/MetaMorph version 7.8 (Molecular Devices, Japan), Adobe Photoshop CS3 v10.0.1 (Adobe, Japan), and NIH ImageJ (https://imagej.nih.gov/ij/). Finally, all chromosomes were counted in at least 10 mitotic cells per line.

### Genomic *in situ* hybridization (GISH) and fluorescence *in situ* hybridization (FISH) analyses of mitotic and meiotic chromosomes

Total genomic DNA was extracted from the young leaves of three lines, 5-7-32 (2*n* = 24), YB2019-30 (2*n* = 25), and YB2019-3 (2*n* = 26), using a DNAs-ici!-R DNA extraction kit (Rizo Inc., Tsukuba, Japan). Extracted DNA was then stored at -30°C until further use. GISH probes were prepared from DNA samples using a Biotin-Nick Translation Mix or a DIG-Nick Translation Mix (both from Sigma, USA). For the rDNA probe, a pTa71 plasmid [40] was also used for DIG-Nick labeling. The 2*n* = 24 probe and rDNA probe were detected using Dig–rhodamine, and the 2*n* = 25 and 2*n* = 26 probes were detected using biotin– streptavidin FITC. All GISH and FISH analyses were conducted using a previously described protocol [41].

### Microdissection and preparation of FISH probes using dissected DNA

Microdissection was performed according to a modified protocol described in a previous report [42]. Briefly, for microdissection chromosome slides prepared from Y-T16-69 (2B) were fist stained with Giemsa solution. Fine glass needles were prepared using a puller (NARISHIGE PC-10) and sterilized using a UV irradiation system (CL-1000, UVP, FUNAKOSHI). Microdissection was then performed using a glass needle attached to a micromanipulator (Eppendorf TransferMan NK 2) under a light microscope (OLYMPUS IX71). Specifically, a single chromosome was microdissected, scraped, then transferred to a PCR tube. The dissected single chromosome—we used one chromosome (i.e., ∼0.37 pg) per tube—was subjected to amplification using a PicoPLEX® Gold Single Cell DNA-Seq Kit (Takara Bio) and indexing primers (DNA HT Dual Index kit, Takara Bio) with all procedures performed as per the manufacturer’s protocol. Four replicates (i.e., #9, #8, #5, and #2) were obtained by repeating the above operation. Next, four DNA probes were prepared using the “Library Amplification” protocol of the PicoPLEX® Gold Single Cell DNA-Seq Kit (50 μl of total volume per reaction; 12 amplification reaction cycles) using 1 μl of 1 mM digoxigenin-11-dUTP (Roche) and 1 μl of purified library + 19 μl of Milli-Q water instead of 20 μl preamplification cleanup DNA. These four DNA probes were then used as FISH probes. FISH was performed as per a previously described protocol [41].

### Estimation of pollen fertility

Mature pollen grains were stained with acetocarmine and classified under a BX43 microscope as fertile (i.e., they showed strong and uniform staining) or sterile (i.e., they showed poor or no staining). Pollen was collected from at least three flowers of KY17-60 (2*n* = 24), KY17-93 (2*n* = 24), KY17-10 (2*n* = 25), KY17- 15 (2*n* = 25), and YB2019-3 (2*n* = 26), and more than 300 pollen grains per line were scored.

### Generation of F_1_ seedlings

Ten female lines (i.e., KY17-5, KY17-6, KY17-7, KY17-21, KY17-29, KY17-30, KY17-38, KY17-43, KY17-45, and Y-T16-69) and seven hermaphroditic lines (i.e., KY17-15, KY17-22, KY17-37, KY17-60, KY17-118, KY17-148, and YB2019-3) were used as parents to produce F_1_ seedlings. Rhizomes of each strain were first divided into 30 g pieces and planted in 15 cm pots in December 2020. Plantlets were grown under natural conditions in Ami-machi, Inashiki-gun, Ibaraki Prefecture (35°99′ N, 140° 20′ E), Japan, until August 2021. Thereafter all plants were moved to a growth chamber and grown for 4 weeks under 16 h light at 25°C/8 h dark at 20°C. During the next 24 days, crosses were performed between the strains with mature pistils and stamens. The cross combinations used are described in detail in subsequent paragraphs. After artificial pollination, plants were grown under the same conditions for another four weeks under 12-h light at 20°C/12-h dark at 15°C and under 12-h light at 15°C/12-h dark at 5°C for a further two weeks. After this point the F_1_ seeds were harvested, sown, then grown in a greenhouse at 25°C.

### Statistical analysis of transmission rates

We calculated the transmission ratio (*k_B_*)—i.e., the mean number of Bs in the microspores or progeny divided by the total number of Bs in the parents—and Z values for parents with B chromosome(s) using a formula described in a previous report [43]. The data were used to determine the accumulation of Bs by comparing *k_B_* with the expected Mendelian rate (0.5). We did not collect data for “embryos”, as in previous studies of other species [43], due to technical difficulties related to embryo observation in *A. lancea*. Instead, we replaced the number of embryos with the number of seeds. Thus, in this study the transmission rates to the next generation are values obtained by compiling meiosis, gametogenesis, reproduction, and seed development. The Z-values indicate significant B accumulation when >1.96 in absolute value, and significant B elimination when Z values <−1.96 [43].

## Results

### Novel extra chromosome(s) in *A. lancea*

Line Y-T16-69 showed typical *A. lancea* morphology (**Fig 1a**). However, its chromosome number was 2*n* = 26 (**Fig 1b**), whereas *A. lancea* reportedly has 2*n* = 24 [34, 35]. Moreover, the number of B chromosomes was stable with no variation between the somatic cells of root tips (**Fig 1b**) and anthers. Therefore, we counted the chromosomes of the 54 lines of *A. lancea* (**S1 Table**) and detected three karyotypes, viz., 2*n* = 24, 2*n* = 25, 2*n* = 26 (**Figs 2a–c** and **Table 1**). Bright DAPI-stained chromosomes were detected in addition to the normal karyotype 2*n* = 24 (**Figs 2a– c**). Thus, 2*n* = 24 is a normal karyotype, while 2*n* = 25 and 2*n* = 26 represent genotypes with 1 and 2 extra chromosomes, respectively. We detected the extra chromosomes in both hermaphroditic and female plants (**Table 1** and **S1 Table**), suggesting a lack of contribution to sex determination. Hereafter, the extra chromosome is referred to as the B chromosome.

**Fig 1.**
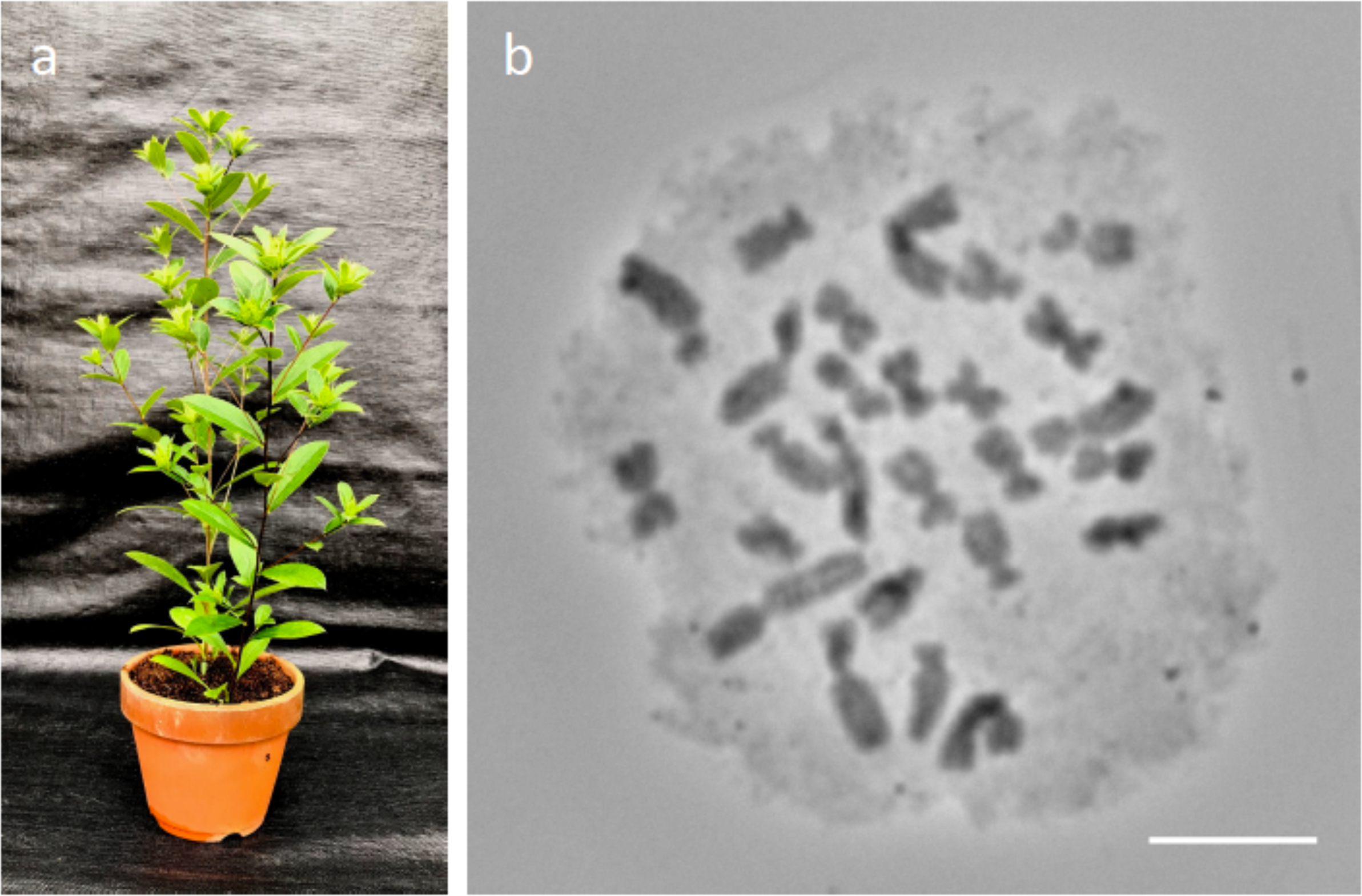
A clone of the *Atractylodes lancea* line Y-T16-69. (a) Morphology typical of the species. (b) Mitotic chromosomes from a root tip (2*n* = 26). Scale bar = 10 µm

**Fig 2.**
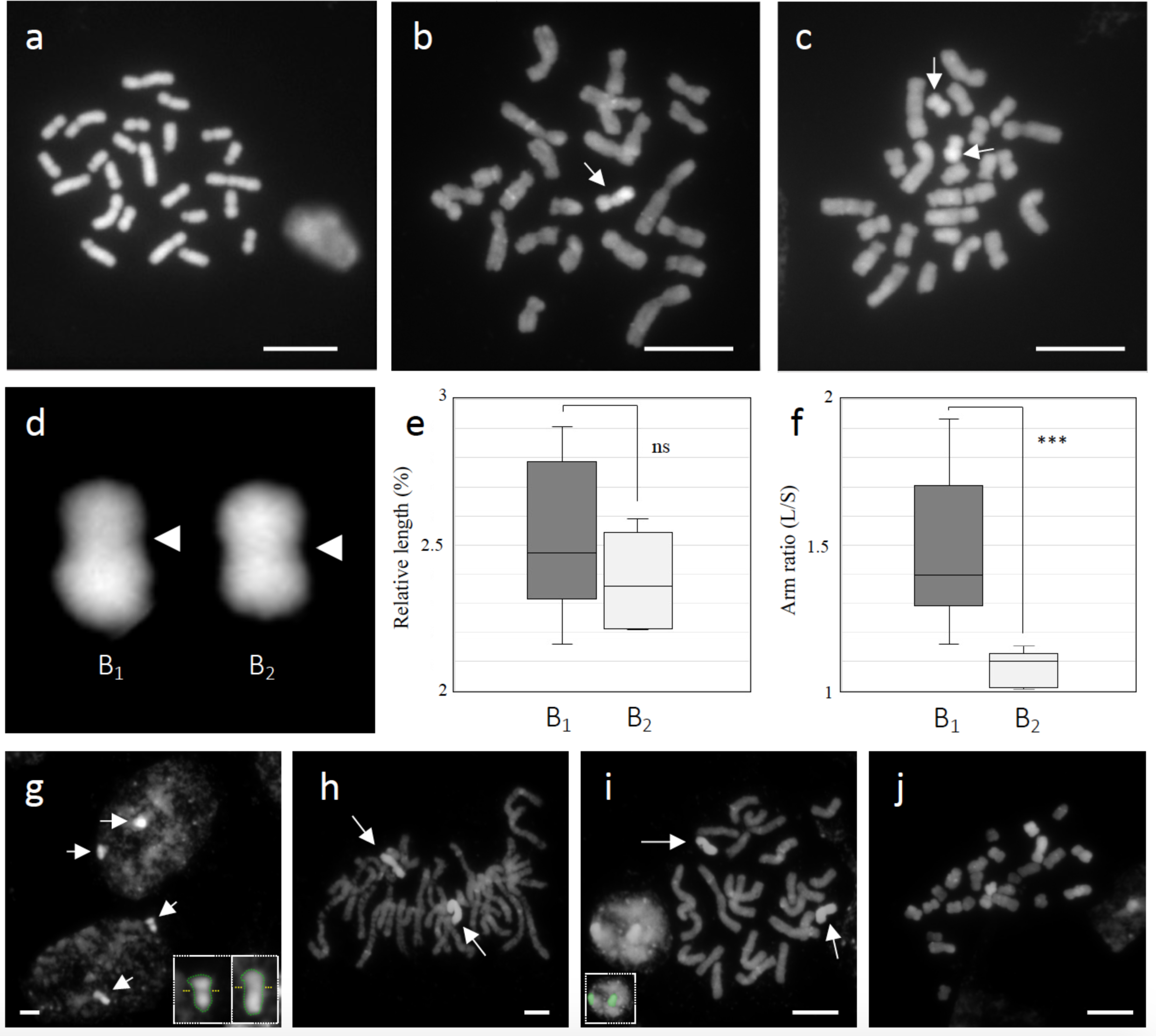
B chromosomes in *A. lancea*. (a) KY17-43 showed 2*n* = 24. (b) KY17-6 (2*n* = 25) showed 24 chromosomes and a DAPI-bright B chromosome (arrow). (c) Y-T16-69 (2*n* = 26) had 24 chromosomes and two B chromosomes (arrows). (d) Cut-out images of two types of B chromosome. Shown are: B_1_, which has distinguishable short and long arms and a metacentric-like B_2_. Arrowheads indicate centromere positions. (e) Y-axes present the relative length (%) of the B chromosome to all 26 chromosomes (i.e., length of the B chromosome (mm) / length of all 26 chromosomes (µm) × 100). No significant differences were detected in the relative length of the two Bs (Student’s *t* test; *P* > 0.1). (f) Y-axes represent arm ratio (L/S: long arm / short arm). A statistically significant difference (*P* < 0.01) was detected in the arm ratios of B_1_ and B_2_. (g–i) Bs (arrows) of Y-T16-69 during the cell cycle. (g) Interphase. DAPI-bright regions of the two Bs can be observed. Note: The size of the DAPI-bright region is comparable to the size of the B chromosome, and often primary constriction can also be observed. Inset: Green dot lines and yellow dot lines indicate the shape of the B chromosome in interphase and primary constriction sites, respectively. (h) Prometaphase. Bs have completed chromatin condensation. (i) Prometaphase. Chromatin condensation of A chromosomes is almost complete. Bs become less visible. Note: The nucleus on the left is more condensed than the nuclei in Fig 2g, and DAPI signals can be observed that indicated the shape of the two B chromosomes (see green colored boxes in inset). (j) Metaphase. Completely condensed chromatin due to anti-tubulin treatment with 2 mM 8-hydroxyquinoline. Further chromosome condensation has made the B chromosomes even less distinguishable from the A chromosomes. Scale bars = 10 µm

**Table 1.** Chromosome numbers in 54 lines of *A. lancea*.

|  | Obs. | Chromosome number |  |  |
| --- | --- | --- | --- | --- |
| | | $2n = 24$ | $2n = 25$ | $2n = 26$ |
| Female plant | 28 | 15 | 11 | 2 |
| Hermaphroditic plant | 26 | 18 | 7 | 1 |
| Total (%) | 54 | 33 (61) | 18 (33) | 3 (6) |

### Two distinct morphologies of B chromosomes

We observed that the two B chromosomes present in line Y-T16-69 had different morphologies (**Fig 2d**). They showed similar lengths relative to the total length of all 26 chromosomes (average relative length = 2.5% ± 0.4 and 2.3% ± 0.2, n = 10), but had different arm ratios (**Figs 2e and 2f**). The submetacentric-like B chromosome (average arm ratio = 1.5 ± 0.2, n = 10) was named B_1_, and the metacentric-like B chromosome (average arm ratio = 1.1 ± 0.1, n = 10) was named B_2_ (**Figs 2d–f**). A difference in the arm ratio was also confirmed from the chromosome images stained with Giemsa (**S1 Appendix**).

### B chromosomes in the cell cycle

Next, DAPI staining revealed that the B chromatin as a DAPI-bright region during the cell cycle (**Figs 2g– j**); moreover, the primary constriction and shape of the B chromosome were visible in the nucleus (**Figs 2g and 2i**). However, identification of Bs was difficult in the metaphase, since other A chromosomes became condensed to the same level (**Fig 2j**).

### Specific DNA of B chromosomes

To determine whether B chromosomes had DNA sequences that were conserved with the A chromosomes, we conducted GISH analysis using labeled genomic DNA used as probes. For this procedures, 2*n* = 24 (0B) was detected as red, and 2*n* = 25 (+1B) or 2*n* = 26 (+2B) was detected as green (**Figs 3a–c**). Moreover, all A chromosomes were yellow (**Fig 3a**). Our results showed that almost B chromosomes were green (**Figs 3b and 3c**), except for the yellow centromeric regions labeled with 0B and the +1B or +2B GISH probes (**Fig 3d**). We interpreted this result as GISH analysis revealing that B chromosomes accumulate specific DNA sequences, with only the centromeric region having sequences similar to those of A chromosomes. FISH analysis was performed using 35S rDNA as a probe (labeled pTa71 clone, typically visualized as NORs) produced two pairs of four FISH signals (which is consistent with a previous report [35]), and showed no signal on the B chromosomes (**S2 Appendix**).

**Fig 3.**
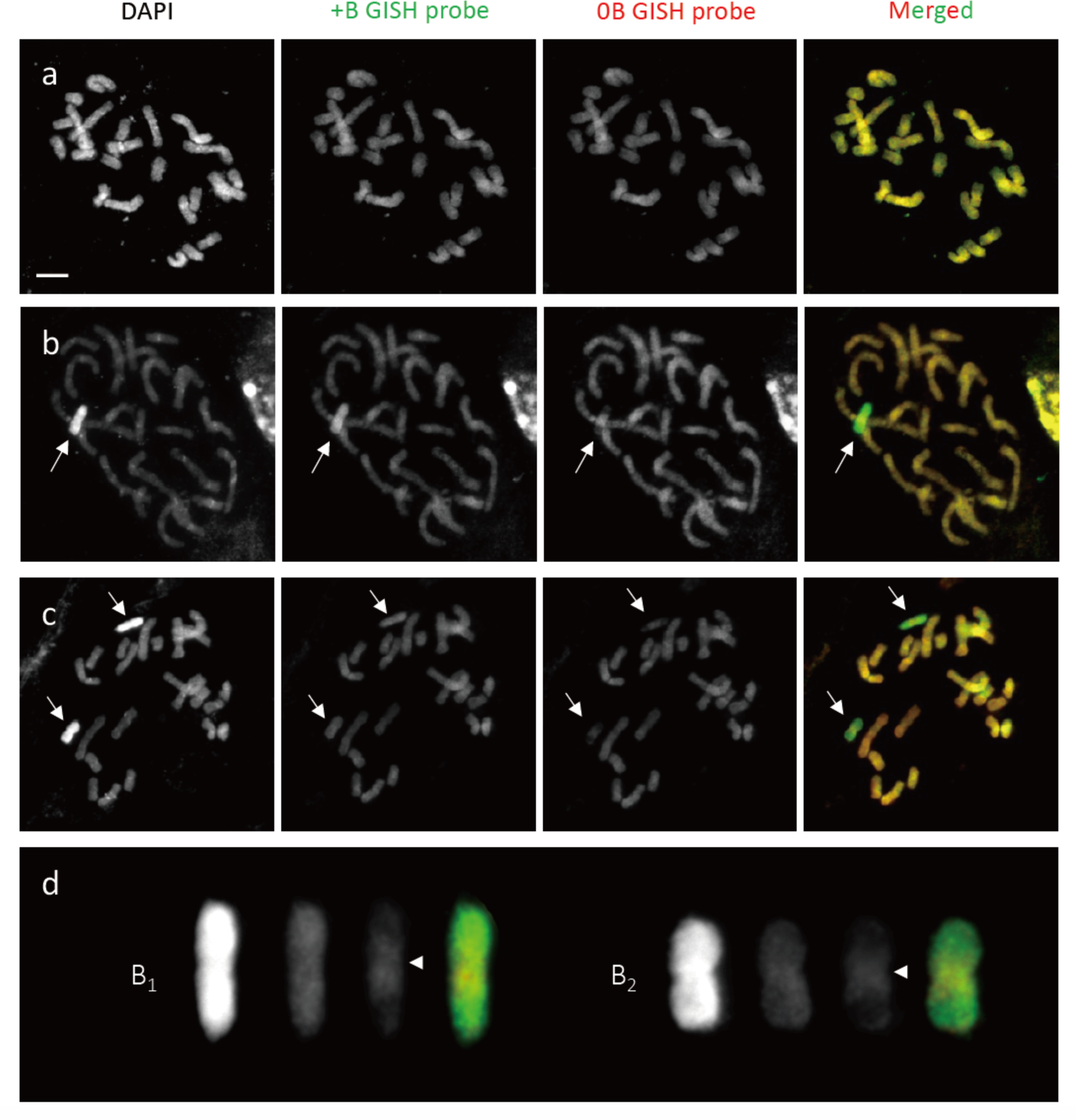
GISH analysis with the gDNA probe (red) for 2*n* = 24 (0B) and gDNA probe (green) for 2*n* = 25 (+1B) or 2*n* = 26 (+2B). (a) 5-7-32 (2*n* = 24). The 24 chromosomes are indicated in yellow in the merged image here and in (b) and (c). (b) YB2019-30 (2*n* = 25). The B chromosome (arrow) is indicated in green. (c) YB2019-3 (2*n* = 26). GISH probe derived from 2*n* =26 gDNA (+2B GISH probe) showing green fluorescence signals on both Bs (arrows). (d) Cut-out images of the Bs shown in panel (c). The centromere regions are hybridized with the 0B probe (arrowheads). Scale bar = 10 µm

### Meiotic behavior of B chromosomes

Next, we observed the behavior of B chromosomes during male meiosis (**Fig 4**). In a 0B plant, 12 bivalents were formed, and regular meiosis was observed without univalent or lagging chromosomes. However, in a 1B plant (KY17-15), a univalent or lagging chromosome appeared from diplotene to anaphase I (**Figs 4a – d and S3 Appendix**). Moreover, since the univalent chromosome can be recognized as a condensed B chromosome in the diplotene stage (**Fig 4a**), we considered the B chromosome as a lagging chromosome. Such abnormal chromosomes did not appear during the second division of meiosis (**Fig 4e**).

**Fig 4.**
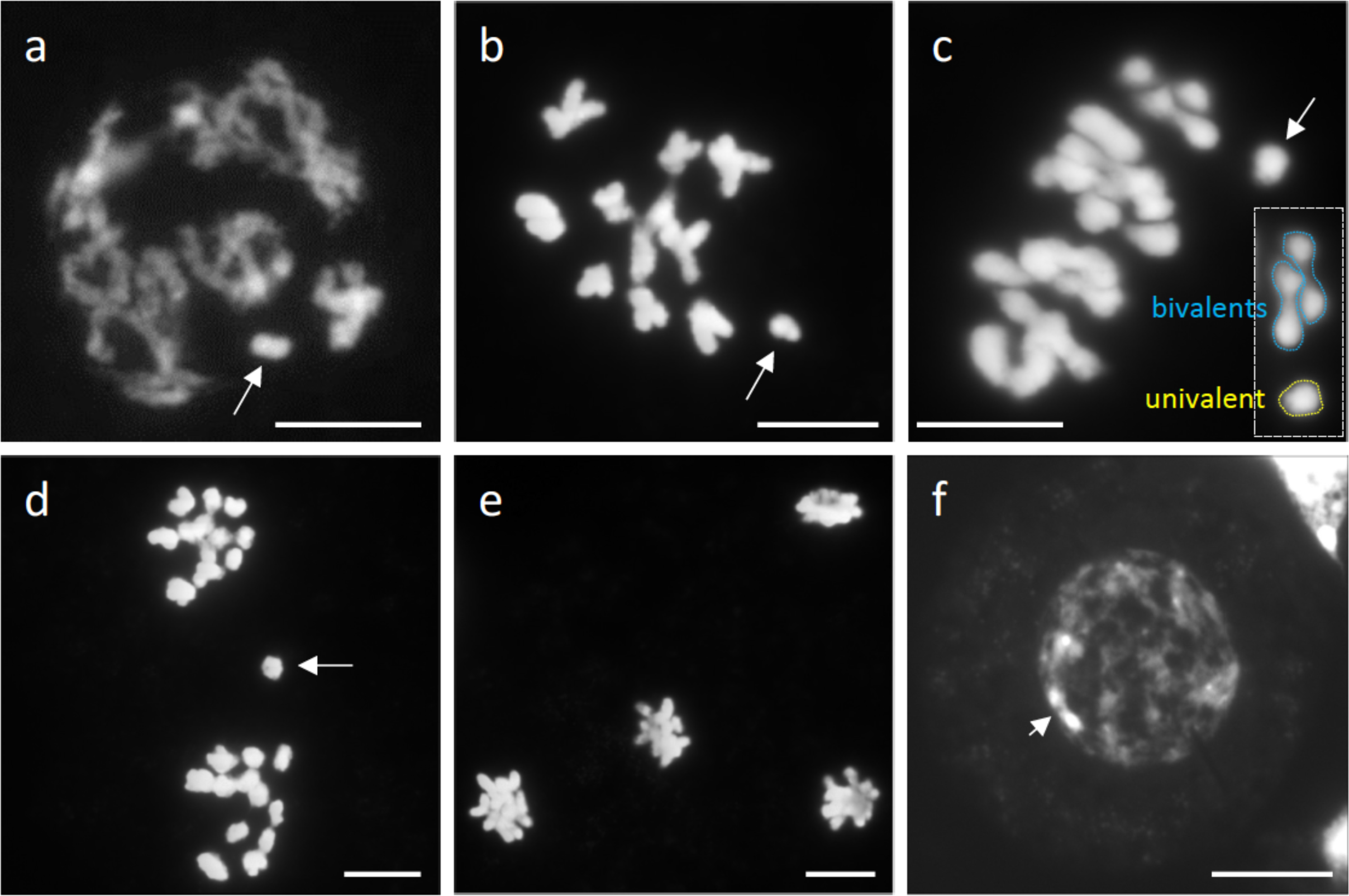
Meiosis in *A. lancea* KY17-15 (2*n* = 25). (a) Diplotene. The condensed B chromosome (arrow) does not pair with other A chromosomes. (b) and (c) Diakinesis and metaphase I. Arrows indicate univalent chromosomes. (d) Telophase I. Arrow indicates a lagging chromosome. (e) Telophase II. No lagging chromosomes or micronuclei are observable. (f) A microspore. The arrow indicates a DAPI-bright region of a B chromosome. Scale bars = 10 µm

### Transition of B chromosomes to microspores

The nuclei of microspores (i.e., the daughter cells produced after male meiosis) showed wither no or only one DAPI-bright signal derived from the B chromosome in 1B plants (**Fig 4f and S4 Appendix**). To confirm that the DAPI-bright region is derived from the B chromosome, we developed B-specific probes by microdissection (**Fig 5**). A total of 11 B chromosomes were excised without distinguishing between B_1_ and B_2_ (**Figs 5a–d**), and the dissected DNAs were amplified for each. For 4 (probes #9, #8, #5, and #2) of the 11 amplified dissected DNAs, FISH analyses of mitotic chromosomes in a 2B line (Y-T16-69) were conducted without any blocking DNAs (**Figs 5e–g and S5 Appendix**). The results indicated that all four probes formed signals on both B chromosomes, suggesting the successful development of the B-specific probes (**Figs 5e–g and S5 Appendix**).

**Fig 5.**
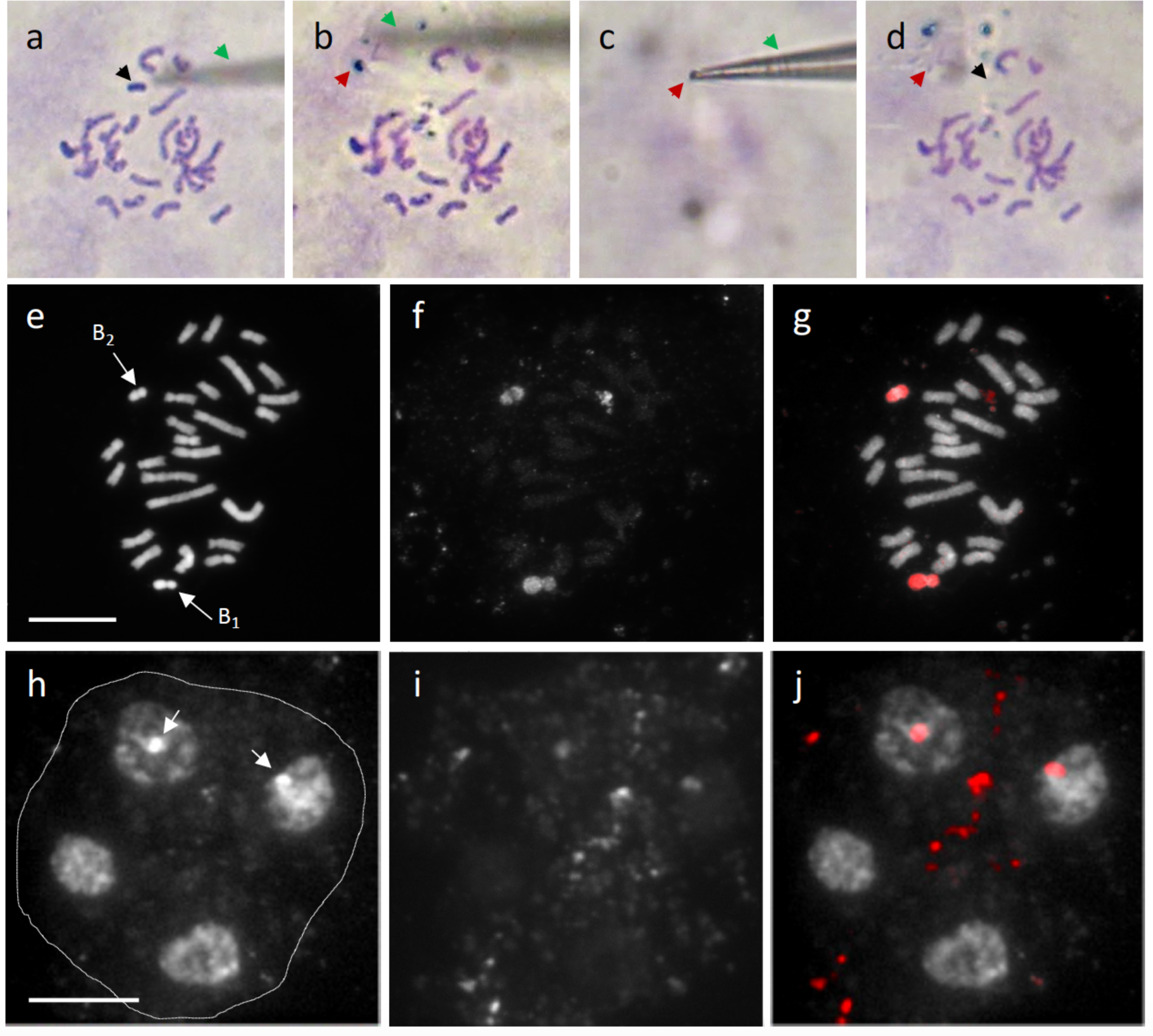
Microdissection of an *A. lanceae* B chromosome. (a–d) Microdissection of an *A. lanceae* B chromosome. (a) A fine glass needle (green arrow) approaches one of the B chromosomes (black arrow). (b) The glass needle (green arrow) scrapes off the target B chromosome. The red arrow indicates a shaved B chromosome. (c) The shaved B chromosome (red arrow) was taken with the tip of a glass needle (green arrow). (d) The spread chromosome following microdissection. Note: Nothing else is present where the target B chromosome (black arrow) and the B chromosome that was scraped off (red arrow) were located. (e–g) FISH probed with microdissected DNA (probe #9). (e) DAPI-stained chromosome spread of Y-T16- 69 carrying B_1_ and B_2_. (f) FISH signals from the dissected DNA probe. (g) Merged image. The dissected DNA probe formed FISH signals on both B chromosomes. (h–j) FISH analysis of a tetrad. (h) DAPI-stained tetrad with four nuclei. The outline of the tetrad is indicated by a line. DAPI-bright regions (arrows) are observed in two of the four nuclei generated by meiosis. (i) FISH signals from the dissected DNA probe (probe #9). (j) Merged image. DAPI-bright regions overlapped with the B-specific probe. Scale bar = 10 µm

Next, using the B-specific probes, we confirmed the B origin of the DAPI-bright region in the nuclei of the tetrad after male meiosis. As a result, the DAPI-bright signals overlapped with B signals (**Figs 5h– j**), indicating that the DAPI-bright region could be counted as the number of B chromosomes. Moreover, the FISH analysis revealed the appearance of micronuclei with B-DNAs (**Fig 6**). In diploids, a single A chromosome is transmitted to the nucleus of each daughter cell by reduction division. We did not observe more than two blocks per nucleus of microspore of 1B line, which would suggest meiotic drive (nondisjunction). Because we observed the micronuclei containing B-DNA in the tetrad, we suspected that the B chromosomes were transmitted lower than Mendelian ratio. Therefore, we investigated the meiotic transmission rate of univalent B chromosomes to microspores using two 1B lines by counting the DAPI- bright region, and found that the B chromosome was transmitted to 18.6%–20.1% of the microspores (**Table 2**), whereas the predicted Mendelian rate was 50%.

**Fig 6.**
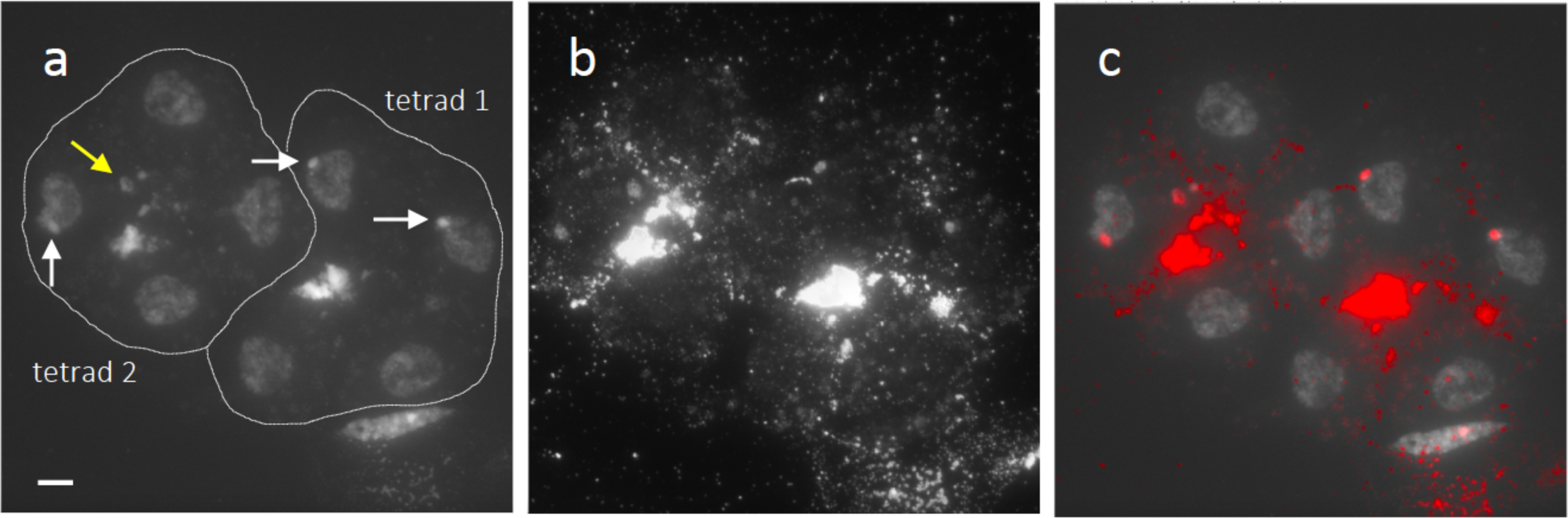
Micronucleus with B-DNAs during male meiosis. (a) Two DAPI-stained tetrads after male meiosis. (b) FISH signal for B-probe #9. (c) Merged image. White arrows and lines indicate DAPI-bright regions with B-DNA and the outline of the tetrad, respectively. The right tetrad (tetrad 1) shows B segregation into two nuclei. The left tetrad (tetrad 2) has only one nucleus carrying B-DNAs, and also has a micronucleus with B-DNAs (yellow arrow). Scale bar = 10 μm.

**Table 2.**
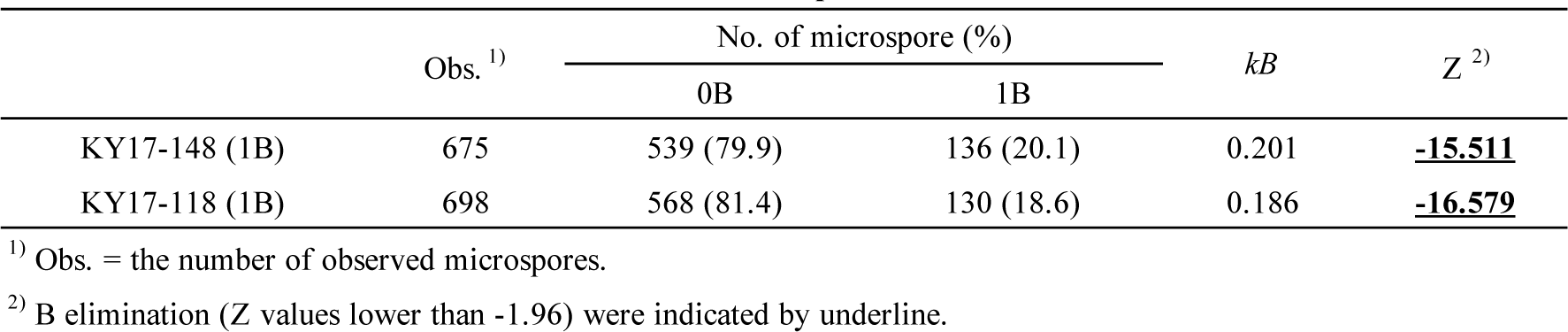
Transmission rate of one B chromosome to microspore.

### Fertility of pollen containing B chromosome

To determine pollen fertility of the 0B, 1B, and 2B plants, the pollen was stained with acetocarmine (**S6 Appendix**). The proportions of well-stained pollen (mature pollen grains) were 98.4% ± 0.7% (n = 3458) for 0B plants, 96.3% ± 0.1% (n = 2728) for 1B plants, and 96.0% ± 0.7% (n = 2131) for 2B plants. Pollen grains of *A. lancea* are trinucleate pollen, and therefore contains three haploid nuclei (i.e., each contains one pollen tube nucleus and two sperm nuclei). No morphological abnormalities among these nuclei were detected in the pollen of 1B plants compared to that of 0B plants (**S6 Appendix**).

### Male transmission rates of B chromosomes

In the 1B plants, the B chromosome is transmitted to 18.6%–20.1% of the microspores. Moreover, pollen containing B chromosome were found not to show morphological abnormalities. We therefore conducted several cross combinations to verify whether pollen with B could be transmitted to subsequent generations. We here collected data on the transmission rate of B chromosomes to the next generation by counting the DAPI-bright regions derived from the B chromosome (**Table 3**, **Figs 7a–f, S2 Table, and S3 Table**). Prior to this analysis, the B-specific probes were used to confirm that the DAPI-bright regions were derived from the B chromosomes in the 1B line, KY17-19 (**Figs 7a–c and S1 Table**).

**Fig 7.**
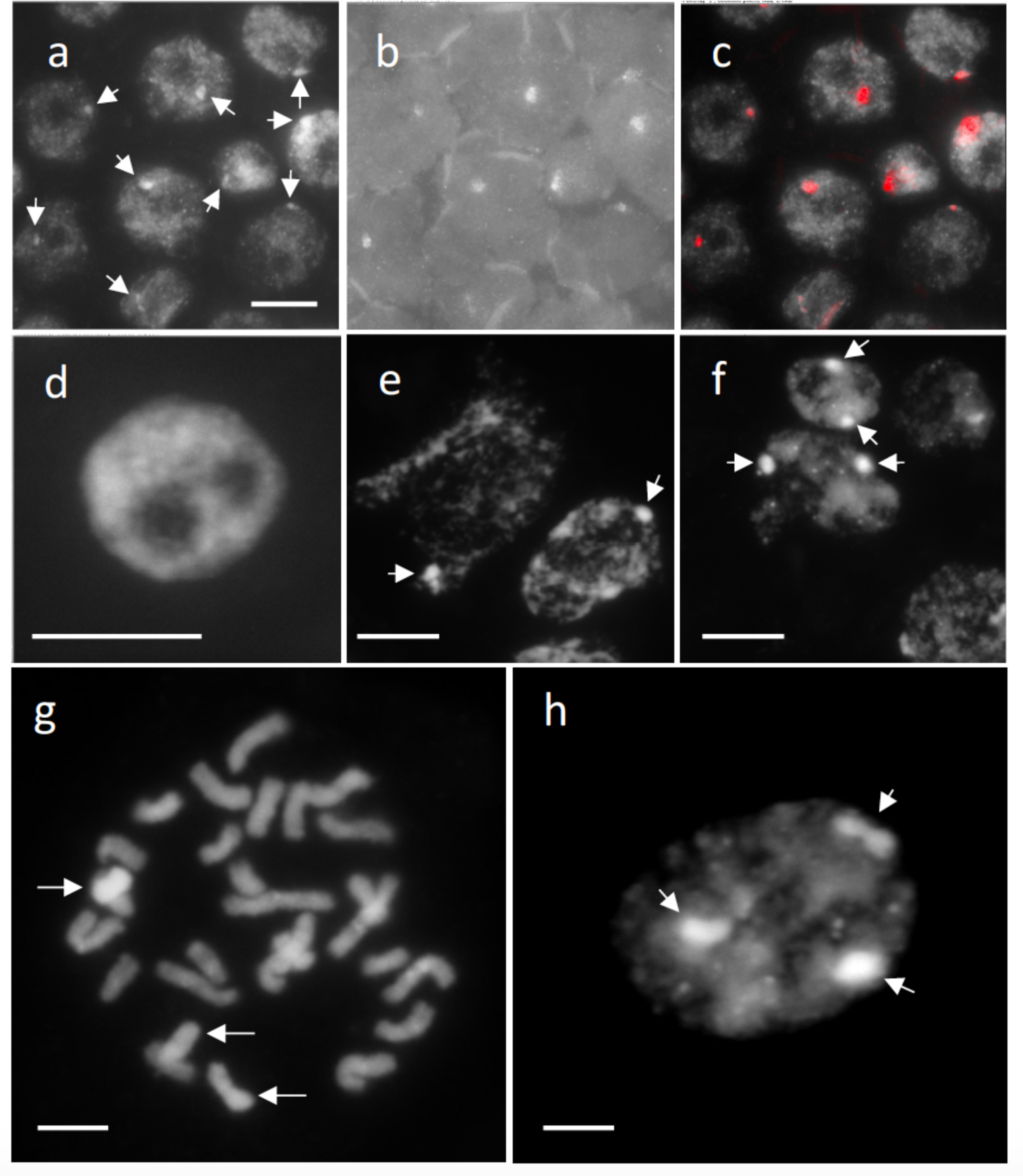
Transmission of B chromosomes to progeny. (a–c) FISH with B probe (probe #9) to the somatic cells of the progeny from a KY17-19 cross (1B; 2*n* = 25. See S1 Table). (a) DAPI-stained nuclei. Arrows indicate B-DAPI-bright regions. (b) FISH signal for the B probe. (c) Merged image. DAPI-bright signals overlapped with the B-specific signals. (d) Nucleus without B-DAPI-bright regions (0B). (e) Nuclei carrying a B- DAPI-bright region (arrows) (1B). (f) Nuclei carrying two B DAPI-bright regions (arrows) (2B). (g–h) Progeny (2*n* = 24 + 3Bs) from the KY17-29 (2*n* = 26) × KY17-22 (2*n* = 24) cross. (a) Mitotic chromosomes. Arrows indicate three Bs. (b) Nucleus. Arrows indicate three DAPI-bright regions derived from three Bs. Scale bar = 10 µm

**Table 3.** Number of B chromosome in progenies from 14 cross combinations.

| Cross combination (female × male) | | Obs. <sup>1)</sup> | No. of progenies (%) | | | | $k_B$ | $Z$ <sup>2)</sup> |
| --- | --- | --- | --- | --- | --- | --- | --- | --- |
|  |  |  | 0B | 1B | 2B | 3B |  |  |
| 0B × 1B | KY17-21 × KY17-148 | 250 | 148 (59.2%) | 102 (40.8%) | - | - | 0.408 | **<u>-2.909</u>** |
|  | KY17-5 × KY17-15 | 33 | 21 (63.6%) | 12 (36.4%) | - | - | 0.364 | -1.567 |
|  | KY17-45 × KY17-118 | 53 | 28 (52.8%) | 25 (47.2%) | - | - | 0.472 | -0.412 |
|  | KY17-45 × KY17-148 | 52 | 29 (55.8%) | 23 (44.2%) | - | - | 0.442 | -0.832 |
| 0B × 2B | KY17-43 × YB2019-3 | 47 | 6 (12.8%) | 29 (61.7%) | 12 (25.5%) | - | 0.564 | 0.875 |
|  | KY17-5 × YB2019-3 | 98 | 23 (23.5%) | 60 (61.2%) | 15 (15.3%) | - | 0.459 | -0.808 |
| 1B × 0B | KY17-7 × KY17-60 | 11 | 2 (18.2%) | 9 (81.8%) | - | - | 0.818 | **<u>2.111</u>** |
|  | KY17-30 × KY17-37 | 10 | 3 (30.0%) | 7 (70.0%) | - | - | 0.700 | 1.265 |
|  | KY17-6 × KY17-22 | 81 | 6 (7.4%) | 75 (92.6%) | - | - | 0.926 | **<u>7.667</u>** |
|  | KY17-38 × KY17-22 | 98 | 35 (35.7%) | 63 (64.3%) | - | - | 0.643 | **<u>2.828</u>** |
| 2B × 0B | KY17-29 × KY17-22 | 77 | 14 (18.2%) | 24 (31.2%) | 38 (49.4%) | 1 (1.2%) | 0.669 | **<u>2.963</u>** |
|  | YT16-69 × KY17-60 | 14 | 0 (0.0%) | 7 (50.0%) | 7 (50.0%) | - | 0.750 | 1.871 |
| 1B × 2B | KY17-30 × YB2019-3 | 51 | 3 (5.9%) | 31 (60.8%) | 17 (33.3%) | 0 (0.0%) | 0.425 | -1.074 |
| 2B × 1B | YT16-69 × KY17-148 | 122 | 5 (4.1%) | 40 (32.8%) | 60 (49.2%) | 17 (13.9%) | 0.577 | 0.003 |

<sup>1)</sup> Number of observed progenies.
<sup>2)</sup> B accumulation (Z values higher than 1.96) and B elimination (Z values lower than -1.96) were indicated by underline.

In the cross of 0B females and 1B males, 36.4%–47.2% of the progeny contained a B chromosome (1B), and the remaining 63.6%–52.8% did not (0B). One (KY17-148) of three males analyzed demonstrated B elimination (*k_B_* = 0.408; Z = –2.909) in a cross with a KY17-21 female; it also showed Mendelian transmission (*k_B_* = 0.442; Z = –0.832) in a cross with a KY17-45 female. Two other males (KY17-15 and KY17-118) also demonstrated Mendelian transmission (**Table 3**). Moreover, the transmission rates (40.8%– 44.2% and 47.2%, respectively) of B chromosomes from KY17-148 (1B) and KY17-118 (1B) males were more than twice those (i.e., 18.6%–20.1%) of microspores containing B chromosomes in these lines (**Table 2**). Finally, a cross involving two 0B females (KY17-43 and KY17-5) showed that one 2B male (YB2019- 3) transmitted its B chromosome at the Mendelian ratio (**Table 3**).

### Female transmission rates of B chromosomes

In three of four cross combinations involving four 1B females and three 0B males, B chromosomes were transmitted to 64.3%–92.6% of all progeny, which suggests B chromosome accumulation (*k_B_* = 0.818, 0.926, and 0.643 and Z = 2.111, 7.667, and 2.828, respectively) (**Table 3**). The remaining cross combination with KY17-30 demonstrated Mendelian transmission (*k_B_* = 0.700; Z = 1.265), although this result may be related to the small number of progenies analyzed (**Table 3**). For the cross between the 2B female (KY17-29 and YT16-69) and 0B male, two 2B females exhibited B chromosome accumulation, but *k_B_* was not significantly different from 0.5 in the cross between YT16-69 and KY17-60 (**Table 3**). We also found one plant (1.2% of the total) from the cross between KY17-29 (2B) and KY17-22 (0B) that showed 2*n* = 27 (3B) in the cross between KY17-29 (2B) and KY17-22 (0B) (**Table 3 and Figs 7g–h**). Moreover, 14 progenies (18.2%) from the cross combination did not carry Bs (**Table 3**); if the two Bs were paired and segregated during meiosis, 0B would not appear.

### Transmission of B chromosomes in 2022 and 2023 and in crossings between B- parents

Next, we tested whether the B transmission rates affected by environmental, a cross between KY17-21 (0B) and KY17-148 (1B) and a cross between KY17-6 (1B) and KY17-60 (0B) were conducted in both 2022 and 2023. We found that the cross between KY17-21 (0B) and KY17-148 (1B) demonstrated B chromosome elimination in 2022 but showed a Mendelian ratio in 2023 (**S2 Table**). In contrast, the cross between KY17-6 (1B) and KY17-60 (0B) demonstrated B chromosome accumulation in both 2022 and 2023 (**S2 Table**).

Finally, to investigate the B transmission rates between B chromosome-containing parents, we performed the 1B × 2B and 2B × 1B crosses. For the 1B × 2B and 2B × 1B crosses, Bs were transmitted in Mendelian ratios (**Table 3**). In addition, similar to that in the cross between KY17-29 (2B) and KY17-22 (0B), we also found 0B progenies (5.9% in 1B × 2B and 4.1% in 2B × 1B) in following crosses (**Table 3**).

## Discussion

### Structure of *A. lancea* B chromosomes

Polymorphisms in the number and morphology of chromosomes can arise from events such as polyploidy, aneuploidy, and chromosomal rearrangements. The extra chromosomes of *A. lancea* detected in this study were determined to be novel B chromosomes because they exhibited no obvious effect on plant morphology and vitality (**Fig 1**), did not pair with an A chromosome (**Fig 4**), and displayed non-Mendelian inheritance from female parents (**Table 3**). We considered the possibility that they are a sex chromosome present in this gynodioecious plant, but we found that the extra chromosome was not a direct sex determinant (**Table 1 and S1 Table**).

Our FISH analyses also indicated that the B chromosomes of *A. lancea* did not contain rDNA (**S2 Appendix**). Based on the B-specific sequences and the sharing of similar DNA in the centromeric region as revealed by in the GISH analysis (**Fig 3**), we inferred two possible evolutionary scenarios, *viz.*, intraspecific construction and hybrid origin. In general, when neo-Bs appear intraspecifically, they can undergo a rapid structural modification that inhibits meiotic pairing with homologous progenitors. Among the various models of B chromosome origin, a report suggests that B chromosomes evolve due to the amplification of B-specific repeats from centromeres or small chromosome fragments [44]. For example, a fragment of the *A. lancea* centromere that shares a sequence with an A chromosome contain rapidly accumulated B-specific repeats to favor stable meiosis and drive, leading to the formation of the present B chromosome. Although we did not determine the DNA sequences specific to *A. lancea* B chromosomes in this study, the constitutive DAPI-bright region resembling heterochromatin suggests the presence of repetitive sequences throughout the chromosome arm. Furthermore, the number of B chromosomes was same in tissue samples taken from both the root tips and anthers of *A. lancea*, and Bs were observed to be stable during somatic cell division. Hence, the *A. lancea* centromere, which acts as the primary constriction and sequence conservation, may also stabilize Bs during somatic cell division. Moreover, this drive may involve newly acquired B sequences rather than differences in the centromere. In rye, the distal region of the long arm of the B chromosome was found to be required for nondisjunction [13]. In the second scenario, Bs are formed from the A chromosomes of closely related species [45]. This hybrid origin of Bs can be detected by the presence of B repeats from closely related species, as has been found in the wasp *Nasonia* [46]. Moreover, the growth areas of the closely related species *Atractylodes chinensis* and *Atractylodes japonica* [47] overlap with that of *A. lancea*, and hybrids between these species exist in natural populations. Nevertheless, there is a lack of sequence information for *A. lancea* Bs, and no evidence from the *A. chinensis* and *A. japonica* karyotypes supports the hybrid-origin scenario.

Sequencing of Bs is a definitive approach to understanding their structure and evolution. Microdissection and cloning of the Bs of *Brachycome dichromosomatica* revealed that the micro B chromosome consists of two tandem repeats (i.e., Bdm29 and Bdm54) that are not present in the A chromosome, indicating that macro B chromosome was not established by simple A chromosome excision [24]. In rye, the B chromosome was isolated by chromosome flow sorting; its shotgun sequencing revealed that the Bs were primarily derived from the rye chromosomes 3R and 7R [48]. In another study, the sequence of the 570-Mbp *Ae. speltoides* B chromosome was determined by comparative sequence analysis and B microdissection [22]. In still another, a high-quality DNA sequence of maize Bs derived from a combination of chromosome flow sorting, Illumina sequencing, Bionano, and Hi-C analyses was determined, and the assembly revealed that the maize B is gene-rich due to continuous transfer from the A chromosome [25]. In this study, we found that the Bs of *A. lancea* are relatively large and display DAPI- bright signals that clearly distinguish them from A chromosomes. Interestingly, this B chromosome could be observed as a DAPI-bright region in the nucleus, and consistently shows the same size and distinct primary constriction. This may indicate that this B remains condensed in the nucleus or is preferentially stained with DAPI to a greater degree than A chromosomes. These structural features helped in conducting microdissection in this study and may be advantageous for chromosome flow sorting, which is required for sequencing analysis.

### Non-Mendelian inheritance of Bs in *A. lancea*

The nature of B transmission observed in *A. lancea* is summarized in **Fig 8**. During male meiosis, the Bs of *A. lancea* plants with 1B formed a univalent chromosome and were transmitted to 18.6%–20.1% of the microspores (i.e., the daughter cells produced by male meiosis), suggesting the loss of univalent B (**Fig 6**) such as interphase elimination or passive B chromosome loss during chromatid segregation [49, 50] without a drive mechanism. In a report studying programmed elimination in *Ae. speltoides*, B chromosomes with active centromere lagged in anaphase and formed micronuclei [22]. In general, a univalent chromosome is easy to eliminate from meiosis; for instance the transmission rate was reported to be as low as 25% in monosomic wheat series [51, 52]. However, the maize *B-9* univalent can reach to one pole ahead of A chromosomes [53], and this chromosome did not show the regular number of the centromere repeats [54, 55]. Although Bs can maintain or even increase their number in male meiosis in other plant and animal species [56, 57], our result suggest that no such male meiotic drive probably exists for *A. lancea*. Since almost all the microspores became mature pollen (**S6 Appendix**), the B chromosome was expected to be transmitted to 18.6%–20.1% of the pollen and the same proportion of the next generation. However, instead it was transmitted to 40.8%–47.2% of the next generation (**Table 3**). Except for the cross KY17-21 × KY17- 148, the transmission rate from male parents did not significantly differ from Mendelian inheritance (0.5) (**Table 3**), but it was approximately twice that of the rate observed for microspores (i.e., 18.6%-20.1%).

**Fig 8.**
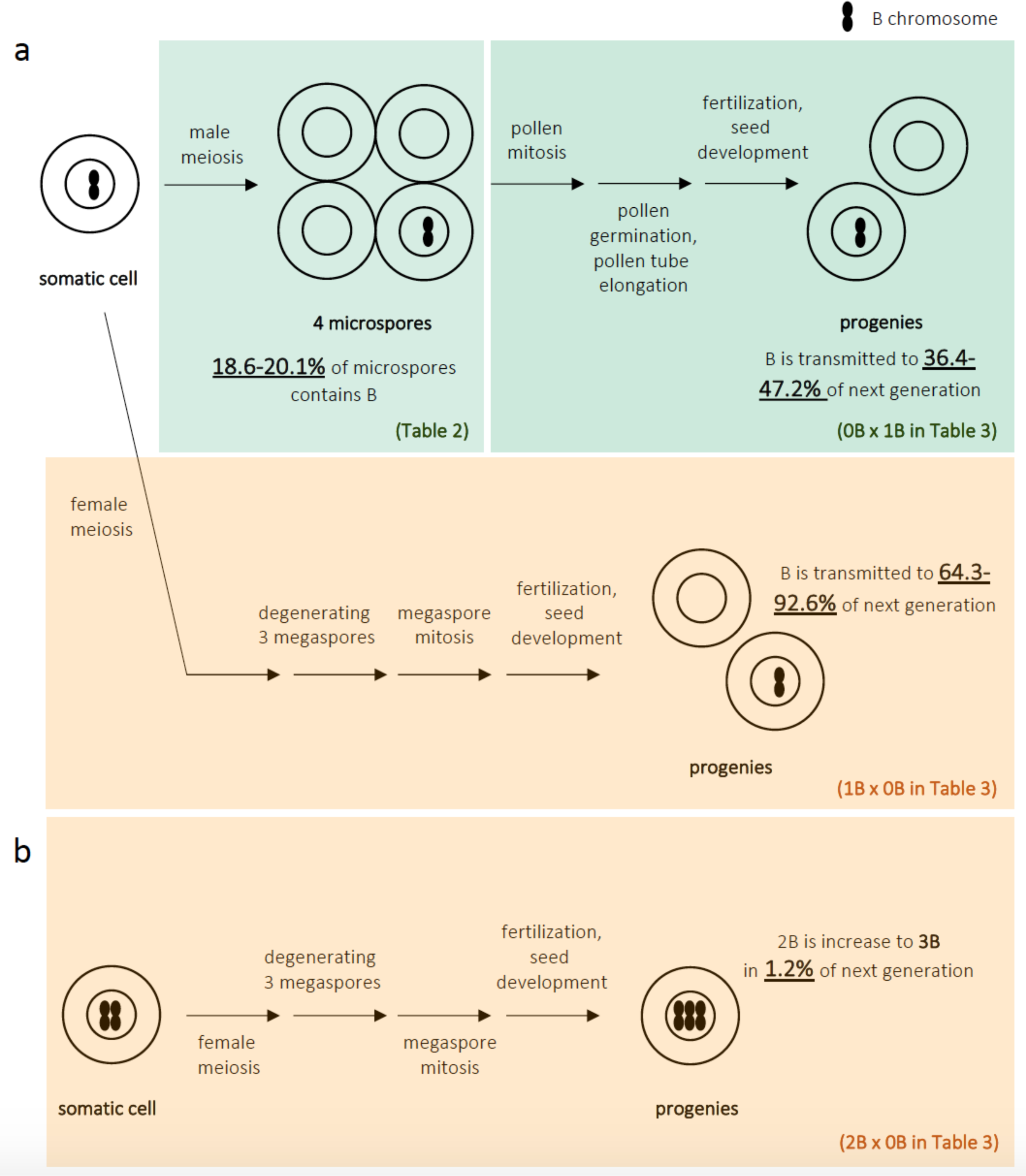
Summary of the transmission of the *A. lancea* B chromosome. (a) In the 1B line, univalent B was transmitted to 18.6%–20.1% of microspores in male meiosis due to chromosome elimination. In contrast, the B chromosome was transmitted to 36.4%–47.2% of cross progeny, suggesting preferential fertilization during pollen mitosis, the germination of pollen grains, or pollen tube growth. In crosses in which the 1B plant is female, the B chromosome was transmitted to 64.3%–92.6% of progeny, suggesting female drive and/or transmission drive during meiosis, megaspore degeneration, fertilization, or seed development. (b) One progeny containing three B chromosomes was found in the 2B × 0B cross, suggesting that nondisjunction occurs during female meiosis, following mitosis (megasporogenesis), fertilization, or seed development.

Moreover, we observed no variation in the number of Bs per cell, as in the case of the nondisjunction of rye and maize B chromosomes. We also detected no abnormalities in the germination of mature seeds or plant growth. Therefore, we speculate that an unknown mechanism involved in preferential reproduction (i.e., fertilization and/or seed development) of pollen carrying Bs may cause a higher rate of B transmission. In rye, *in vitro* tests of pollen grain germination and pollen tube growth show that pollen from plants carrying Bs exhibit better performance than pollen produced from plants without Bs [58]. A previous study has also reported on the effect of Bs on development in grasshopper embryos [59]. The B-specific probes developed in this study may be helpful for further analysis of the mechanistic reason for higher rate of B transmission in pollen mitosis, germination of pollen grains, pollen tube growth, and fertilization in *A. lancea*.

Unlike the transmission from male parents, maternal univalent B chromosomes were transmitted to 64.3%–92.6% of progenies, and B chromosome accumulation was significantly present in at least three of four cross combinations (**Fig 8 and Table 3**). Although we have no cytological evidence, the higher transmission rate of *A. lancea* Bs may be explained by the female drive. This is because the asymmetries in female meiosis and meiotic daughter cell (megaspore) formation provide an opportunity for meiotic drive. One of the four megaspores develops into an egg cell, whereas the remaining three are not involved in reproduction. Bs may cause female meiotic drive and are thereby preferentially transmitted to the nucleus of cells that will become egg cells, resulting in a transmission rate >0.5. During female meiosis in *Lilium callosum*, one univalent B chromosome has been found to lie on the micropylar side of the equatorial plate and is transmitted to 80% of the next generation [21]. Furthermore, in rye and *Ae. speltoides*, extensive observations suggest that the asymmetric formation of microtubules induces nondisjunction rather than a dysfunctional centromere [11, 14, 60]. Interestingly, a recent study of Drosophila by Hanlon and Hawley [61] showed that the female B transmission to progeny in a specific genetic background is non-drive, but in a mutant background the authors observed a biased transmission of the B chromosomes on the female side, thereby indicating a meiotic drive suppression system.

Similar to the 1B × 0B cross, *A. lancea* 2B female plants also tended to preferentially transmit Bs to the next generation (**Table 3**). However, if two B chromosomes pair during meiosis, the B chromosome should be transmitted at a rate of 100%, which is inconsistent with the appearance of 0B progeny (i.e., 18.2% from KY17-29 × KY17-22 cross, 5.9% from KY17-30 × YB2019-3 cross, and 4.1% from YT16-69 × KY17-148 cross). We also found 2*n* = 27 progeny (1.2%) in a cross of 2B × 0B *A. lancea* plants (**Table 3**). The mechanism of B chromosome transmission, including those responsible for the rare cases listed above, needs to be clarified by future cytological analyses. In a cross between 2B and 1B, Bs were transmitted according to the Mendelian ratio, which may be due to progeny of the transmission ratio of B accumulation on the female side and weak B elimination on the male side. (although the transmission ratios from most male parents are not significantly different from Mendelian transmission). Alternatively, since there are no >3B lines were observed among the 54 lines analyzed, the accumulation of the B chromosome might be harmful in *A. lancea*, although the 2B lines used in this study did not show weaknesses relative to 0B and 1B lines.

## Conclusions

In this paper, we report the identification of a new B chromosome in *A. lancea*. Subsequently, we characterized its structure and the biased B transmissions. The Bs were appeared as DAPI-bright region in the nuclei, suggesting an unusual chromatin condensation pattern or preferential staining with DAPI. On the male side, Bs are lost during micronucleation or chromatid segregation, but there seems to be a mechanism by which gametes with Bs are preferentially fertilized. On the female side, a relatively strong drive was found; this transmitted Bs to ∼92.6% of the progeny. The biased B transmission pattern among male and female parents is unique and may provide new insights into B chromosome biology.

## Author contributions

S.K., T.T., H.T., K.S., and S.I. designed the research. K.H., S.K., M.I., T.T., K.S., and S.I. performed the research. T.T., M.S. prepared all materials including F_1_ seedlings; K.H., S.K., M.I., T.T., K.S., and S.I. analyzed the data and K.H., S.K., and T.T. wrote the paper. All authors have read, discussed, and approved the final version of this manuscript.

## Abbreviations

Bs: B chromosomes
FISH: fluorescence *in situ* hybridization
GISH: genomic *in situ* hybridization
NORs: nucleolar organizer regions
rDNA: ribosomal DNA

---

**S1 Appendix.**
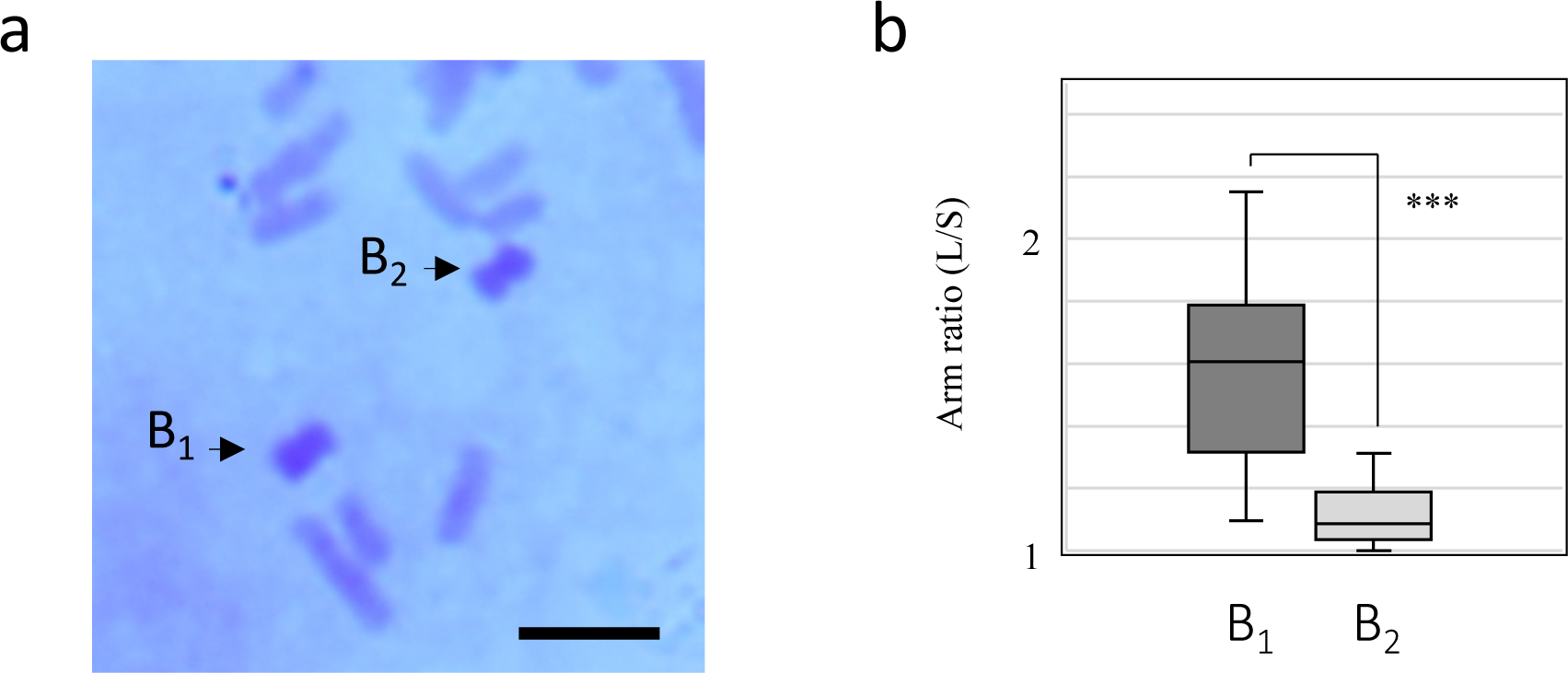
Arm ratios of B_1_ and B_2_ in Giemsa−stained chromosome images. (A) B_1_ and B_2_ stained with Giemsa solution. Scale bar = 10 μm. (b) Y-axes show arm ratios (L/S: long arm / short arm). A statistically significant difference (P < 0.01) was detected for the arm ratios of B_1_ and B_2_.

**S2 Appendix.**
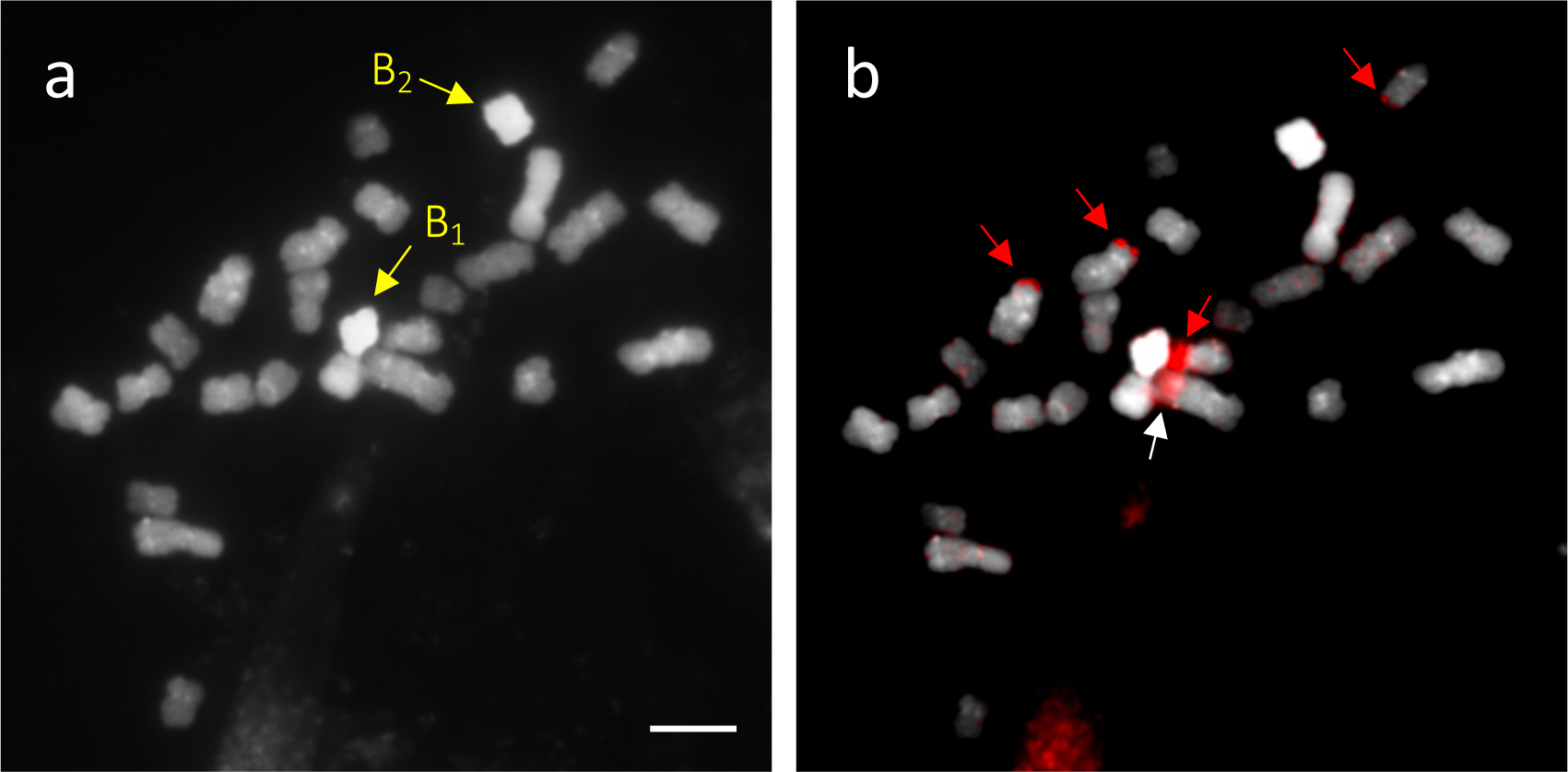
35S rDNA-FISH of *A. lancea* line Y-T16-69 (2*n* = 26, 2B). (a) DAPI-stained mitotic chromosomes. (b) The FISH image. The B chromosomes (i.e., B_1_ and B_2_, indicated by yellow arrows) do not contain rDNA. Red arrows indicate the four loci of 35S rDNA. White arrow indicates a non-specific signal. Scale bar = 10 μm.

**S3 Appendix.**
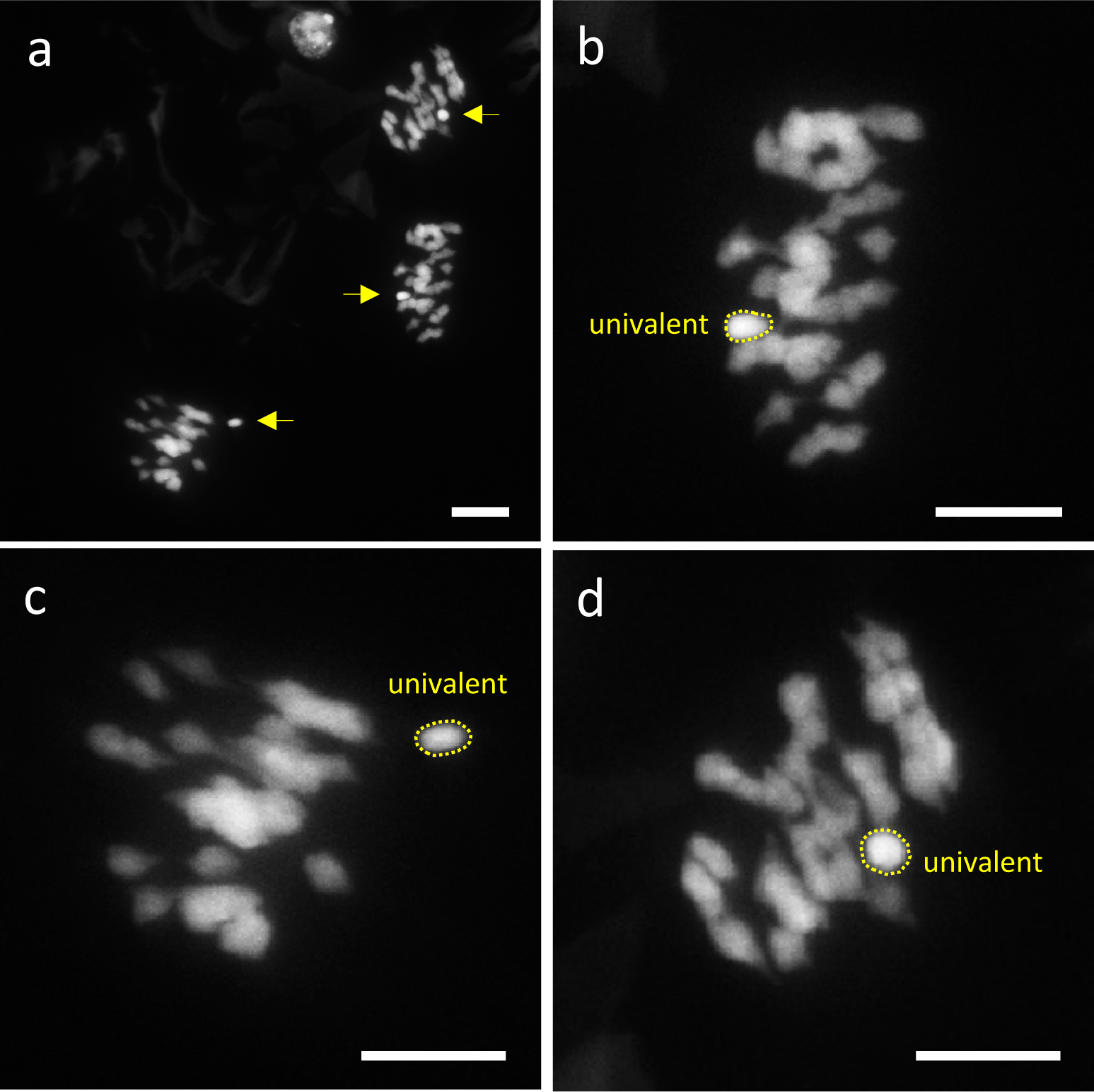
DAPI-bright univalent in meiotic metaphase I. (a) Three DAPI-stained meiotic metaphase I cells. Univalent chromosomes (yellow arrows) are observed in each cell. (b–d) Close-up images of the three metaphase I cells. Scale bars = 10 μm.

**S4 Appendix.**
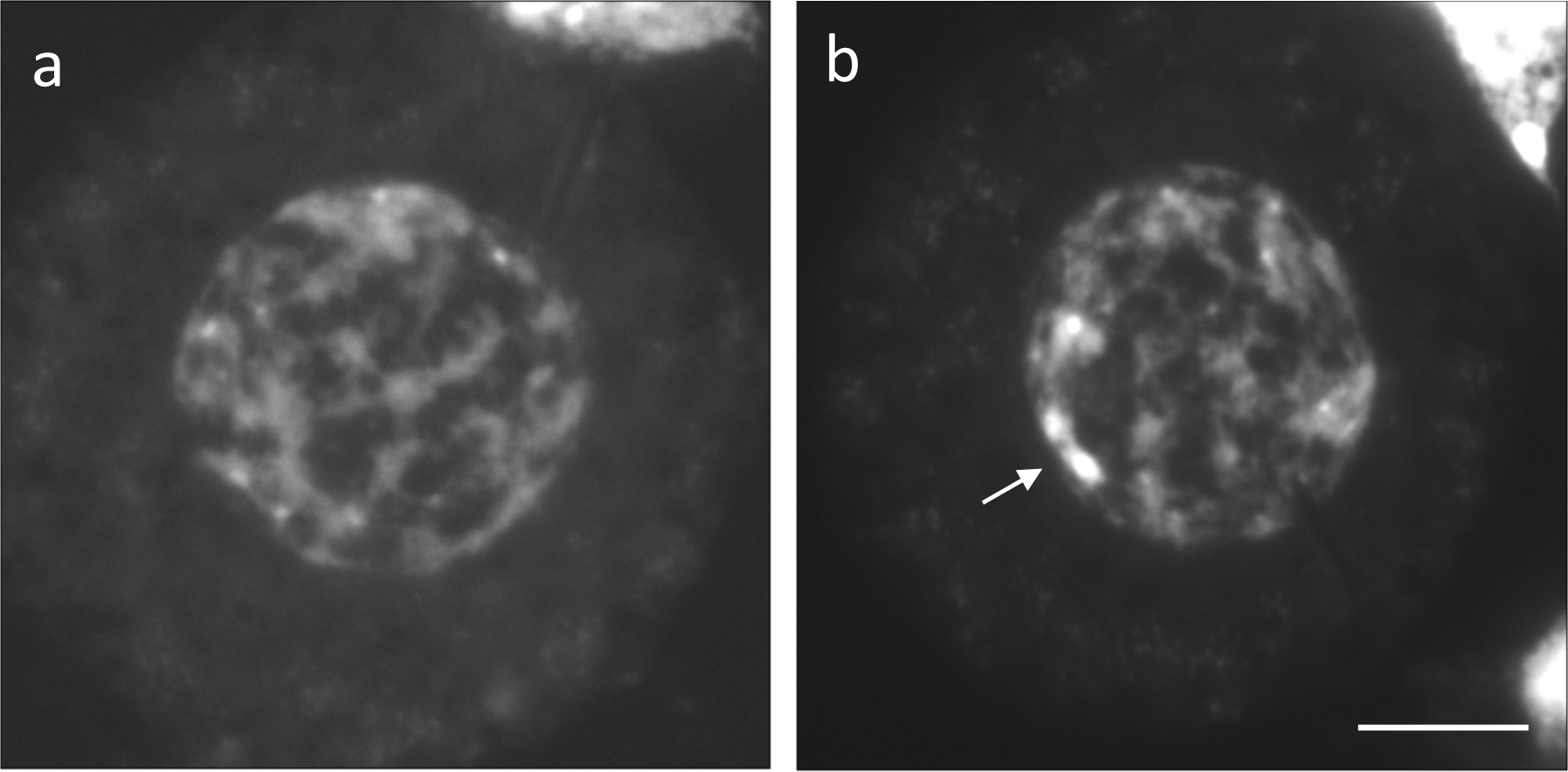
Microspore after male meiosis in KY17-15 (1B). (a) The nucleus of the microspore does not show a DAPI-bright region. (b) Nucleus of the microspore containing a DAPI-bright region. Scale bar = 10 μm.

**S5 Appendix.**
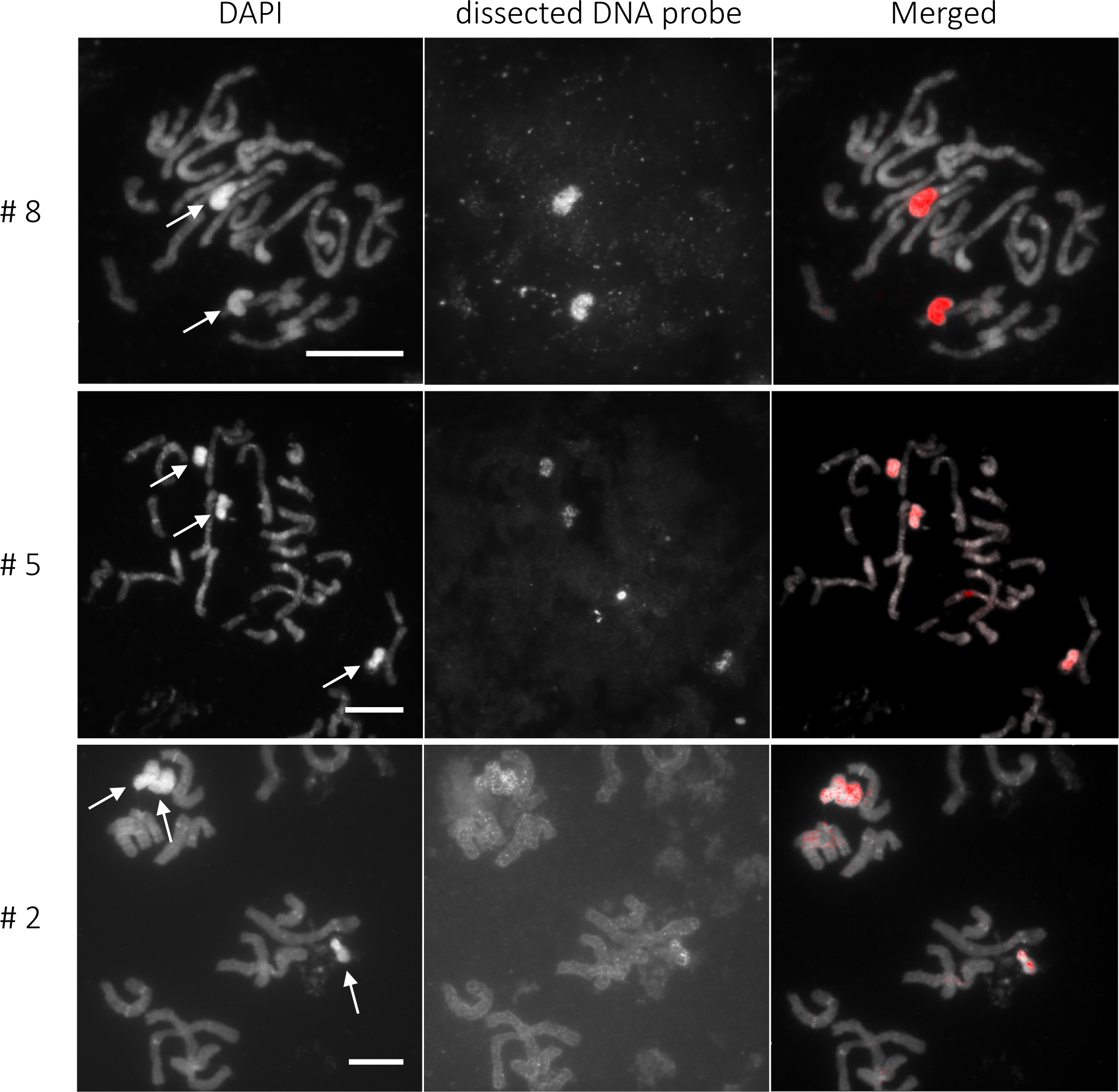
FISH analysis probed with three dissected DNA samples (i.e., #8, 5, 2). All three probes hybridize specifically to B chromosomes (arrows). Scale bars = 10 μm.

**S6 Appendix.**
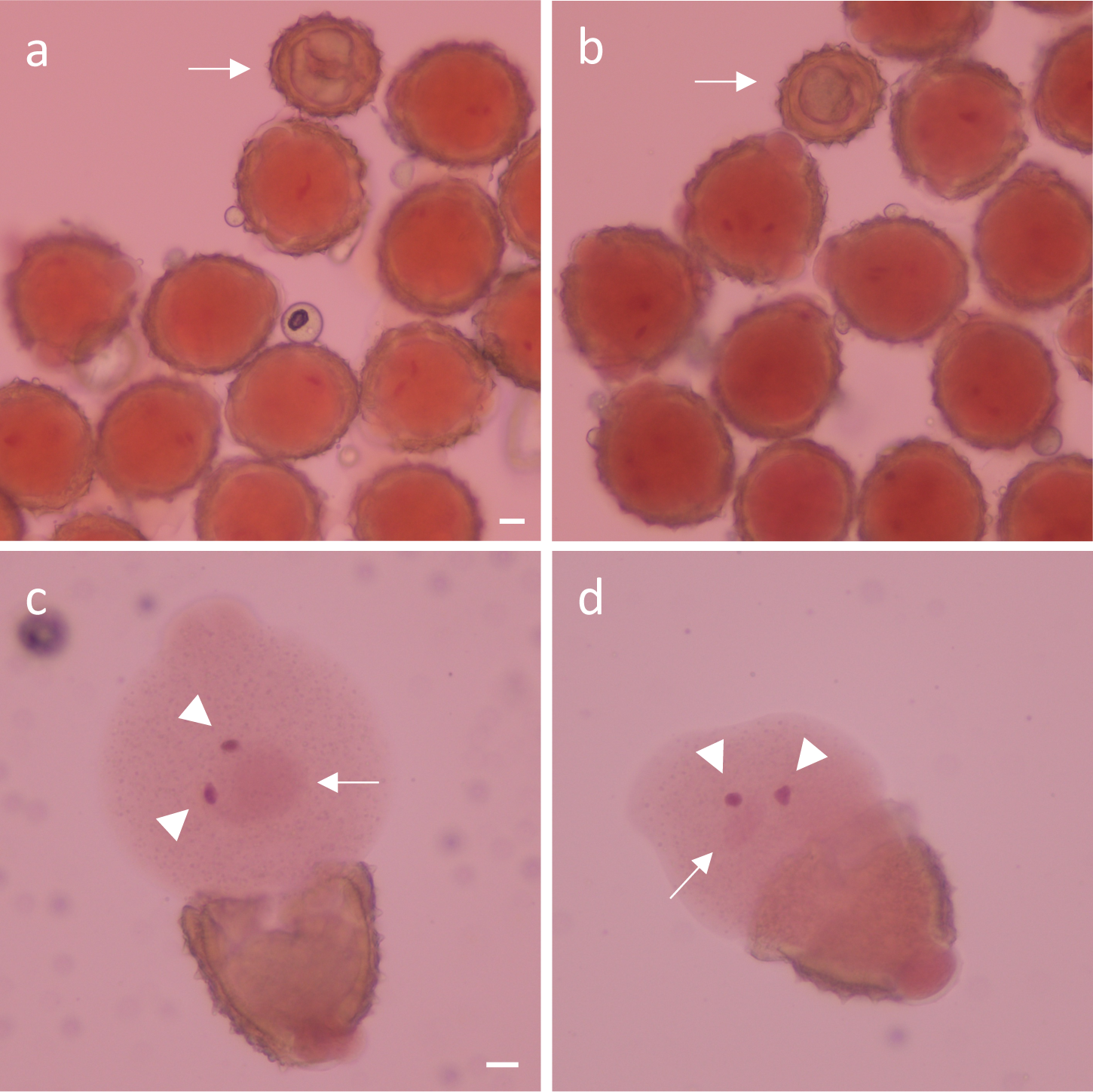
Acetocarmine staining of pollen grains. (a) KY17-60 (0B). The cytoplasm of most pollen grains was stained; the arrow indicates an unstained grain. (b) KY17-15 (1B). (c, d) Two generative nuclei and a vegetative (pollen tube) nucleus (c) KY17-60 (0B) and (d) KY17-15 (1B). No morphological differences were observed between the nuclei in (c) and (d) . Scale bars = 10 µm

**S1 Table.** Chromosome number (2*n*) and B chromosome in 54 *A. lancea* lines.

| Line ID | Sex phenotype | $2n$ | No. of B chr. <sup>1)</sup> |
| --- | --- | --- | --- |
| Y-T16-69 | female | 26 | 2 |
| 5-7-32 | female | 24 | 0 |
| 5-2-12 | female | 24 | 0 |
| 5-12-7 | female | 24 | 0 |
| KY17-4 | female | 24 | 0 |
| KY17-5 | female | 24 | 0 |
| KY17-6 | female | 25 | 1 |
| KY17-7 | female | 25 | 1 |
| KY17-8 | female | 24 | 0 |
| KY17-10 | hermaphroditic | 25 | 1 |
| KY17-15 | hermaphroditic | 25 | 1 |
| KY17-16 | female | 25 | 1 |
| KY17-19 | female | 25 | 1 |
| KY17-21 | female | 24 | 0 |
| KY17-22 | hermaphroditic | 24 | 0 |
| KY17-28 | hermaphroditic | 24 | 0 |
| KY17-29 | female | 26 | 2 |
| KY17-30 | female | 25 | 1 |
| KY17-34 | female | 24 | 0 |
| KY17-37 | hermaphroditic | 24 | 0 |
| KY17-38 | female | 25 | 1 |
| KY17-41 | female | 25 | 1 |
| KY17-43 | female | 24 | 0 |
| KY17-45 | female | 24 | 0 |
| KY17-47 | hermaphroditic | 24 | 0 |
| KY17-48 | hermaphroditic | 24 | 0 |
| KY17-51 | female | 24 | 0 |
| KY17-57 | hermaphroditic | 24 | 0 |
| KY17-59 | female | 25 | 1 |
| KY17-60 | hermaphroditic | 24 | 0 |
| KY17-61 | female | 24 | 0 |
| KY17-62 | female | 25 | 1 |
| KY17-69 | female | 24 | 0 |
| KY17-75 | hermaphroditic | 24 | 0 |
| KY17-84 | female | 25 | 1 |
| KY17-93 | hermaphroditic | 24 | 0 |
| KY17-100 | female | 25 | 1 |
| KY17-113 | female | 24 | 0 |
| KY17-117 | hermaphroditic | 24 | 0 |
| KY17-118 | hermaphroditic | 25 | 1 |
| KY17-122 | female | 24 | 0 |
| KY17-143 | hermaphroditic | 24 | 0 |
| KY17-146 | hermaphroditic | 24 | 0 |
| KY17-148 | hermaphroditic | 25 | 1 |
| KY17-151 | hermaphroditic | 24 | 0 |
| KY17-171 | hermaphroditic | 24 | 0 |
| YB2019-3 | hermaphroditic | 26 | 2 |
| YB2019-5 | hermaphroditic | 24 | 0 |
| YB2019-30 | hermaphroditic | 25 | 1 |
| YB2019-34 | hermaphroditic | 24 | 0 |
| YB2019-35 | hermaphroditic | 25 | 1 |
| YB201937 | hermaphroditic | 24 | 0 |
| YB2019-54 | hermaphroditic | 24 | 0 |
| YB2019-119 | hermaphroditic | 25 | 1 |

<sup>1)</sup> Lines carrying B chromosome(s) are highlighted.

**S2 Table.** Number of B chromosome in progenies from 2 cross combinations in 2022 and 2023.

| Female × Male | Cross combination | Year | Obs. | No. of progenies (%) | | | | $k_B$ | $Z^{1)}$ |
| --- | --- | --- | --- | --- | --- | --- | --- | --- | --- |
|  |  |  |  | 0B | 1B | 2B | 3B |  |  |
| 0 B × 1 B | KY17-21 × KY17-148 | 2022 | 147 | 91 (61.9%) | 56 (38.1%) | - | - | 0.381 | **<u>-2.887</u>** |
|  | KY17-21 × KY17-148 | 2023 | 103 | 57 (55.3%) | 46 (44.7%) | - | - | 0.447 | -1.084 |
| 1 B × 0 B | KY17-6 × KY17-22 | 2022 | 32 | 3 (9.4%) | 29 (90.6%) | - | - | 0.906 | **<u>4.596</u>** |
|  | KY17-6 × KY17-22 | 2023 | 49 | 3 (6.1%) | 46 (93.9%) | - | - | 0.939 | **<u>6.143</u>** |

<sup>1)</sup> B accumulation (Z values higher than 1.96) and B elimination (Z values lower than -1.96) were indicated by underline.

